# Loss of the scavenger receptor MARCO results in uncontrolled vomocytosis of fungi from macrophages

**DOI:** 10.1101/2024.02.22.581301

**Authors:** Chinaemerem U. Onyishi, Gyorgy Fejer, Subhankar Mukhopadhyay, Siamon Gordon, Robin C. May

## Abstract

Vomocytosis, also known as nonlytic exocytosis, is a process whereby fully phagocytosed microbes are expelled from phagocytes without discernible damage to either the phagocyte or microbe. Although this phenomenon was first described in the opportunistic fungal pathogen *Cryptococcus neoformans* in 2006, to date, mechanistic studies have been hampered by an inability to reliably stimulate or inhibit vomocytosis. Here we present the fortuitous discovery that macrophages lacking the scavenger receptor MAcrophage Receptor with COllagenous domain (MARCO), exhibit near-total vomocytosis of internalised cryptococci within a few hours of infection. Our findings suggest that MARCO’s role in modulating vomocytosis is independent of its role as a phagocytic receptor and instead may be driven by variation in cytoskeletal arrangement between wildtype and MARCO-deficient macrophages.

## Introduction

*Cryptococcus neoformans* is an opportunist fungal pathogen that causes life-threatening meningitis, mainly in immunocompromised individuals such as HIV/AIDS patients (1). Infection is thought to begin with the inhalation of the fungi into the lungs where it encounters macrophages of the innate immune system that serve as the first line of defence against infection (1). The interaction between *C. neoformans* and macrophages can lead to a range of outcomes including fungal survival and replication within macrophages (2,3), lateral transfer of cryptococci between macrophages (4,5), lysis of macrophages (6) and vomocytosis, also called nonlytic exocytosis (7,8).

Vomocytosis is a nonlytic expulsion mechanism where fully phagocytosed fungi are expelled from the macrophage with no evidence of host cell damage (7–9). Vomocytosis occurs through the fusion of *Cryptococcus*-containing phagosome with the plasma membrane in a manner that is modulated by the actin cytoskeleton (10). It has also been shown to require phagosome membrane permeabilization (10) and a failure to fully acidify the phagosome (11,12). Previous studies have identified the mitogen-activated protein kinase ERK5 and the phospholipid binding protein Annexin A2 as regulators of vomocytosis, with ERK5 inhibition and Annexin A2 deficiency leading to increased and decreased vomocytosis, respectively (13,14). Moreover, stimulation of macrophages with type 1 interferons (IFNα and IFNβ), mimicking viral coinfection, increased cryptococcal vomocytosis (15). Very little else is known about host regulators of vomocytosis.

Here we present the chance observation that the scavenger receptor MAcrophage Receptor with Collagenous structure (MARCO) is a key modulator of vomocytosis. Typically, vomocytosis rates from wildtype macrophages are between 10-20%, but this number rises close to 100% in *Marco*^*-/-*^ macrophages. Further investigation indicates that this impact on vomocytosis is likely independent of MARCO’s role in phagocytosis but may instead result from the previously documented cytoskeletal dysfunction seen in *Marco*^*-/-*^ macrophages (16). As well as providing a powerful experimental tool for the future investigations into this phenomenon, this finding also has important implications for interpreting infection assays conducted in *Marco*^*-/-*^ animals.

## Results and Discussion

### *Marco*^*-/-*^ macrophages show increased vomocytosis of non-opsonised *C. neoformans*

While investigating the role of scavenger receptors in the phagocytosis of *C. neoformans* using a non-transformed GM-CSF dependent alveolar-like macrophage cell line derived from wildtype and *Marco*^*-/-*^ C57BL/6 mice (17), we observed that, in LPS-stimulated macrophages, MARCO-deficiency led to decreased non-opsonic phagocytosis of *C. neoformans* (Figure 1A). Using live cell imaging to observe the interaction between *C. neoformans* and macrophages, we noted a dramatic increase in the vomocytosis of *C. neoformans* from *Marco*^*-/-*^ macrophages with 80% to 100% of infected macrophages experiencing at least one vomocytosis event (Figure 1B; Supplementary Video 1). No other host or pathogen factor has been found to increase the rate of vomocytosis to such an extent. Not only did *Marco*^*-/-*^ macrophages show elevated vomocytosis, but 75% of nonlytic expulsion events occurred within the first 1 h 40 mins of the initiation of the timelapse video, with the median time to a vomocytosis event being 0.92 h (55 mins) (Figure 1C; Supplementary Video 2). In contrast, vomocytosis in wildtype macrophages occurred over a wider range of time with the median time to vomocytosis being 10.6 h (10 h:35 mins) (Figure 1C). Figure 1D provides a representative image of non-opsonised *C. neoformans* being expelled from *Marco*^*-/-*^ macrophages.

**Figure 1:**
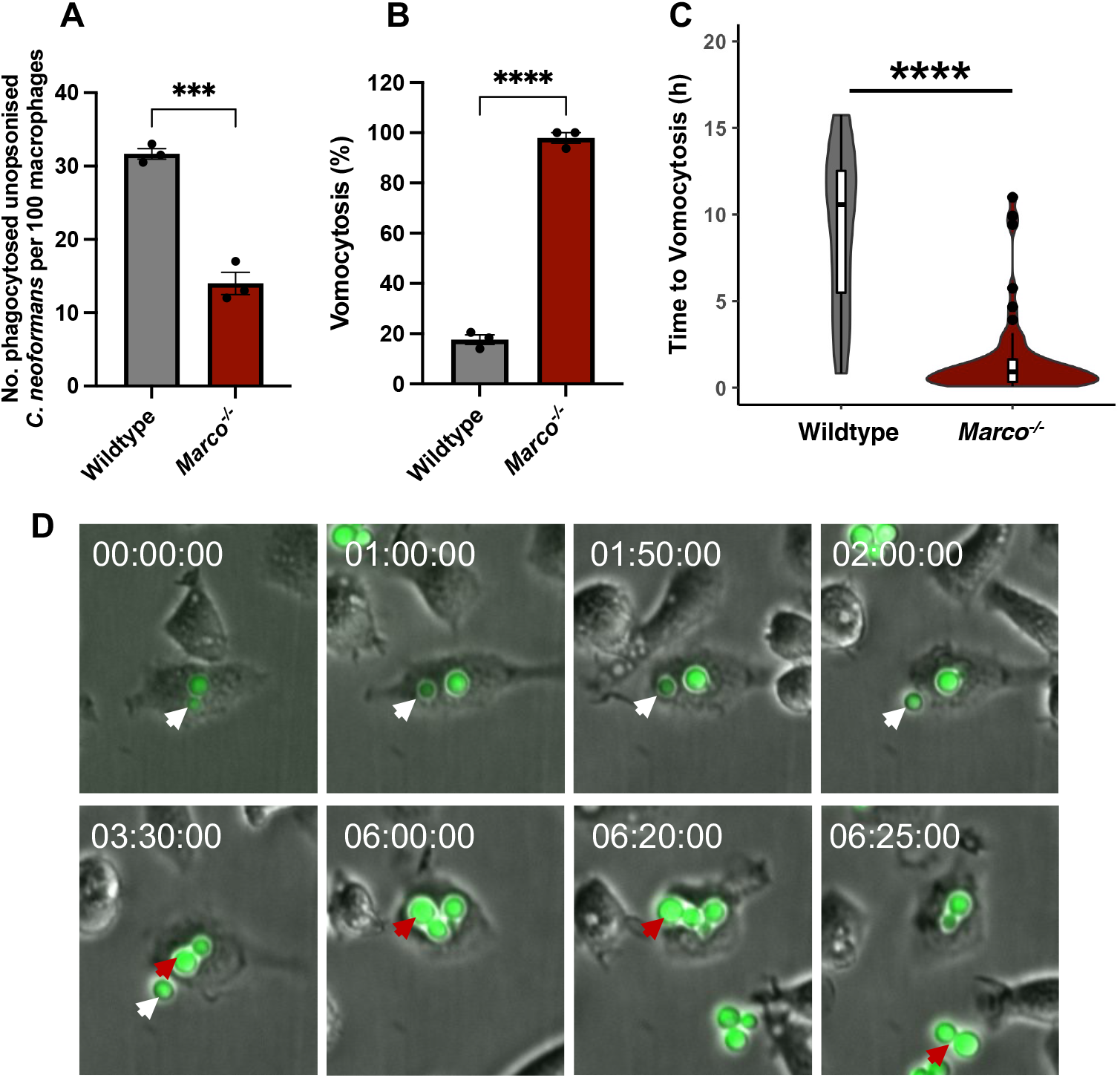
Macrophages were stimulated overnight with LPS then infected with non-opsonised *Cryptococcus neoformans*. **(A)** Phagocytosis was quantified as the number of internalised cryptococci per 100 macrophages. **(B)** Vomocytosis was quantified over a 16 h period and presented as the percentage of infected macrophages that experienced one or more vomocytosis events. At least 200 macrophages were observed. Data is presented as mean ± SEM; ***p<0.001, ****p<0.0001 in an unpaired two sided t-test. **(C)** The time at which individual vomocytosis events took place was quantified and expressed as decimals. Wildtype (n=32); *Marco*^*-/-*^ (n=56). A violin plot with an overlapping box plot was created using the ggplot2 package on R; ****p<0.0001 in a Mann-Whitney test. **(D)** Representative image showing vomocytosis of GFP-expressing *C. neoformans* from *Marco*^*-/-*^ macrophages. Time is presented in hh:mm:ss; white and red arrows follow the course of expulsion events. Data is representative of three independent experiments.

### MARCO-deficiency leads to elevated vomocytosis of 18B7 antibody-opsonised *C. neoformans* and yeast-locked *Candida albicans*, but not heat killed *C. neoformans* or latex beads

To investigate the generality of this phenomena, we infected macrophages with heat-killed *C. neoformans*, 7 μm diameter latex beads, anti-GXM 18B7 antibody-opsonised *C. neoformans*, and a yeast-locked TetOn-NRG1 *C. albicans* strain that constitutively expresses Nrg1 transcription factor, thereby preventing yeast to hypha formation (18). In line with previous data showing that inert particles do not undergo vomocytosis (7,8), we observed no vomocytosis of heat-killed *C. neoformans* by either wildtype or *Marco*^*-/-*^ macrophages out of 55 infected macrophages observed (Table 1), and only a single event (amongst 239 infected cells) when macrophages were “infected” with latex beads (Table 2). Notably, *Marco*^*-/-*^ macrophages showed decreased phagocytosis of heat-killed cryptococci (Supplementary Figure 1A) and latex beads (Supplementary Figure 1B) compared to wildtype cells.

**Table 1:**
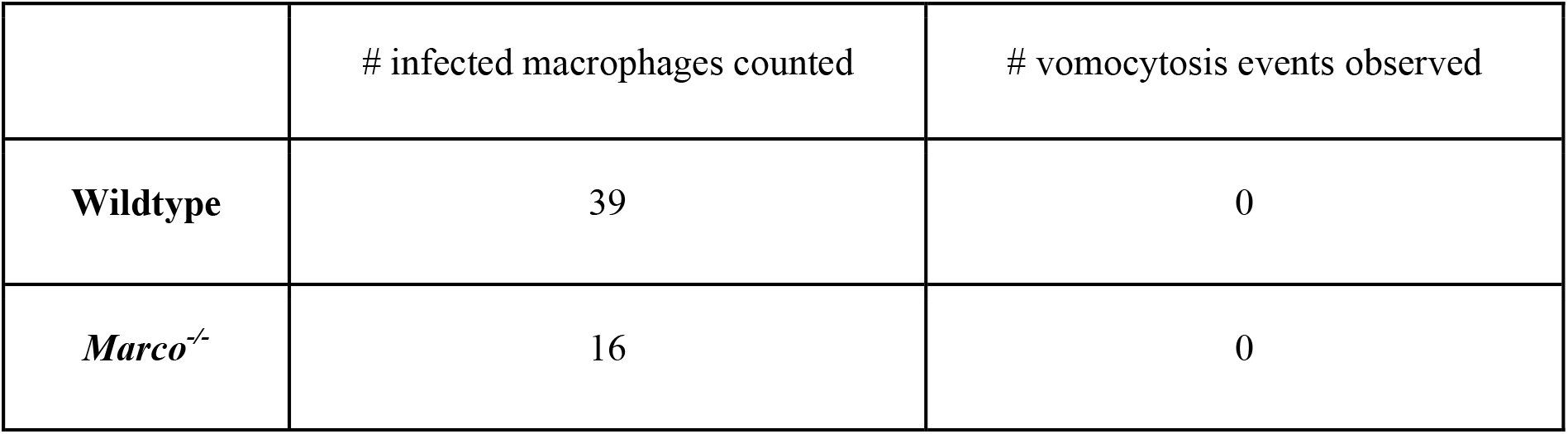
Quantification of the vomocytosis of heat-killed *C. neoformans* in wildtype and *Marco*^*-/-*^ macrophages.

|  | # infected macrophages counted | # vomocytosis events observed |
| --- | --- | --- |
| **Wildtype** | 39 | 0 |
| ***Marco*<sup>-/-</sup>** | 16 | 0 |

**Table 2:**
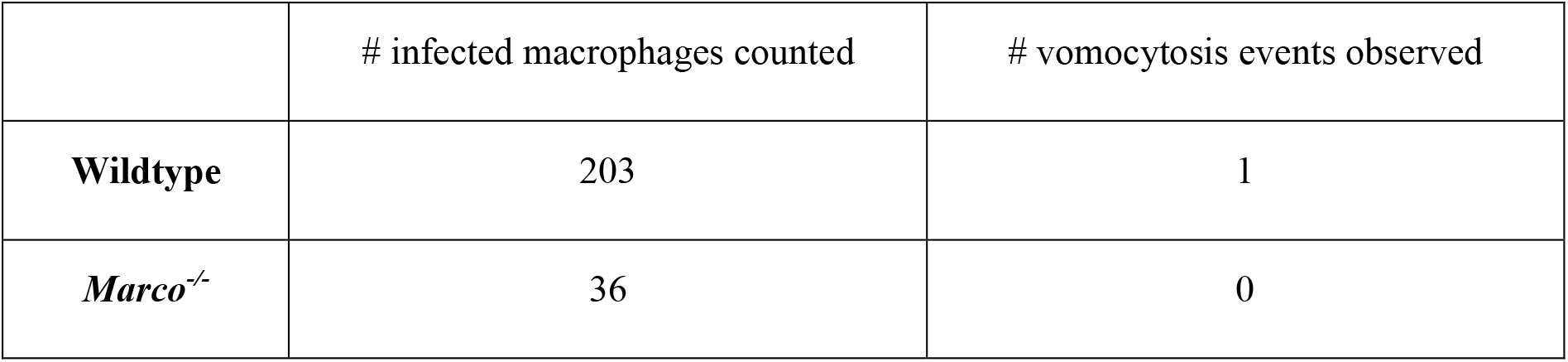
Quantification of the vomocytosis of latex beads in wildtype and *Marco*^*-/-*^ macrophages.

|  | # infected macrophages counted | # vomocytosis events observed |
| --- | --- | --- |
| **Wildtype** | 203 | 1 |
| ***Marco*<sup>-/-</sup>** | 36 | 0 |

Next, macrophages were infected with antibody-opsonised fungi to drive uptake via FcγRs. As expected, there was no difference in antibody-opsonised phagocytosis between wildtype and *Marco*^*-/-*^ macrophages (Figure 2A). However, vomocytosis was elevated in *Marco*^*-/-*^ macrophages compared to wildtype cells (Figure 2B). Finally, macrophages were infected with a yeast-locked *C. albicans* strain that fail to undergo filamentation and observed over a 6 h period. As expected, since phagocytosis of *Candida* is predominantly driven by Dectin-1 (19), there was no difference in phagocytosis between wildtype and *Marco*^*-/-*^ macrophages (Figure 2C). Surprisingly, we observed increased vomocytosis of yeast-locked *Candida* from MARCO-deficient macrophages (Figure 2D and E; Supplementary Video 3). The percentage of *Marco*^*-/-*^ macrophages that experienced at least one vomocytosis event was not as dramatic as that observed with *Cryptococcus*; however, vomocytosis of wildtype *C. albicans* is rare, happening at a rate of <1% over a 6-hour period (20). Therefore, a rate of 30% over 6 hours in *Marco*^*-/-*^ cells is significant for this fungal pathogen. Given that elevated vomocytosis was observed when phagocytosis was mediated by non-opsonic receptors (Figure 1B), FcγR (Figure 2B) and Dectin-1 (Figure 2D), it seems likely that the role of MARCO in vomocytosis is independent of the mechanism of uptake.

**Figure 2:**
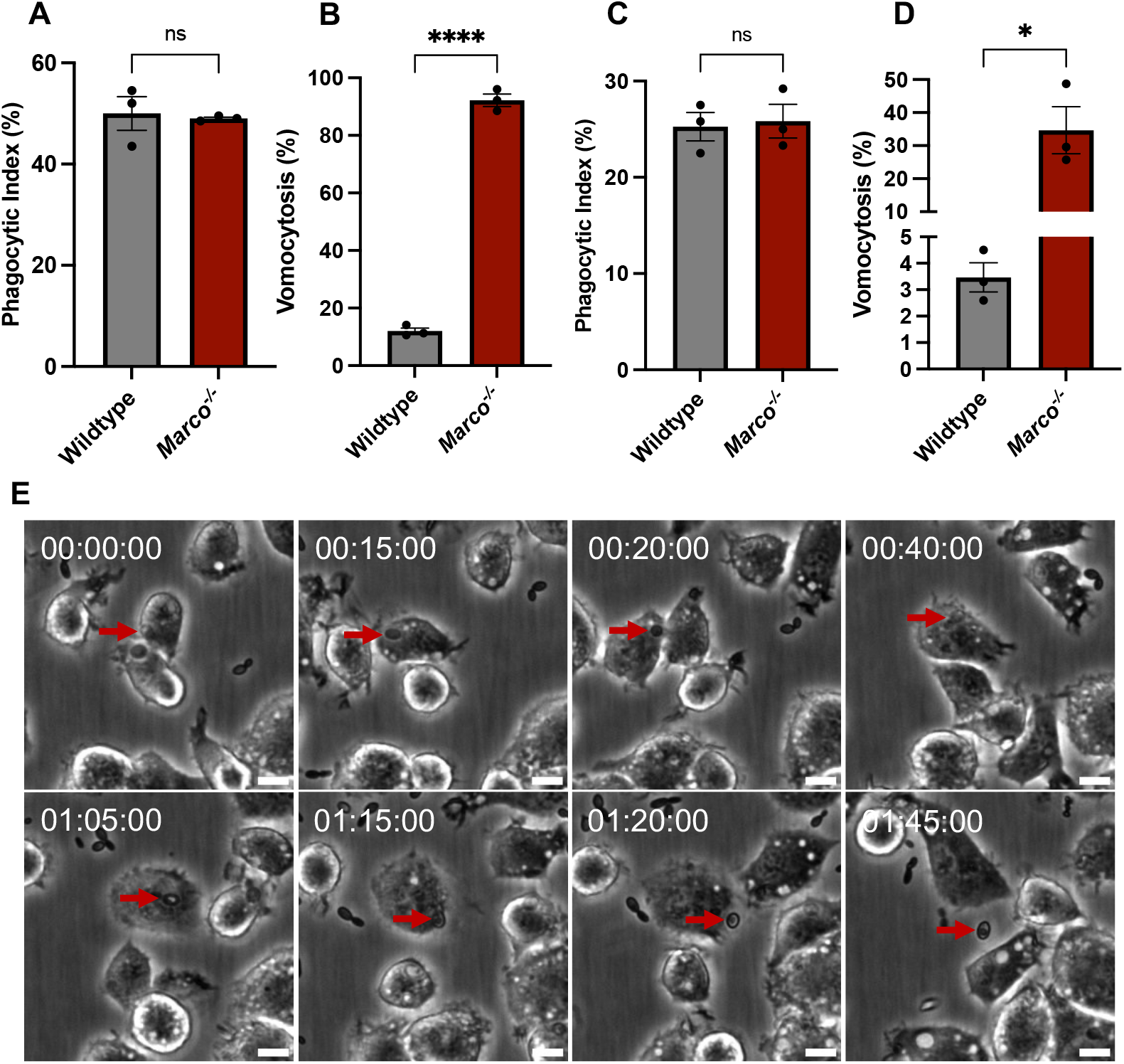
**(A, B)** Wildtype and *Marco*^*-/-*^ macrophages were stimulated with 10 ng/mL LPS overnight, then infected with anti-GXM 18B7 antibody opsonised *C. neoformans*. After 2 h infection, images were acquired every 5 mins for 16 h. **(C, D)** LPS stimulated macrophages were infected with a yeast-locked TetO-NRG1 *C. albicans* strain that constitutively expresses Nrg1 transcription factor, thereby preventing yeast to hyphae transition. Images were acquired every 5 mins for 6 h. **(A, C)** Phagocytic index (%) represents the percentage of macrophages that phagocytosed one or more fungal cells. **(B, D)** Vomocytosis (%) is the percentage of infected macrophages that experienced at least one expulsion events. At least 200 macrophages were observed per condition. Data is representative of two independent experiments. Data shown as mean SEM; ns, not significant; *p<0.05; ****p<0.0001 in a t-test. **(E)** Representative image showing vomocytosis of *C. albicans* from *Marco*^*-/-*^ MPI cells. Time is presented in hh:mm:ss; red arrows follow the course of a vomocytosis event; scale bar = 10 μm.

### Treatment of wildtype MPI cells with inhibitors of MARCO does not phenocopy increased vomocytosis seen in *Marco*^*-/-*^ cells

To explore whether the vomocytosis phenotype seen in *Marco*^*-/-*^ can be induced in wildtype macrophages, we exposed wildtype cells to polyguanylic acid potassium salt (polyG), a MARCO ligand and inhibitor (21–23), and quantified vomocytosis. Although polyG pre-treatment decreased the phagocytosis of non-opsonised *C. neoformans* (Figure 3A), unlike genetic knockout of *MARCO*, the inhibition of MARCO using a ligand did not result in an increase in vomocytosis (Figure 3B). Since polyG functions as a competitive inhibitor and likely does not block MARCO-mediated downstream signalling, this suggests that the impact of MARCO on vomocytosis can be mechanistically separated from its ligand-binding activity.

**Figure 3:**
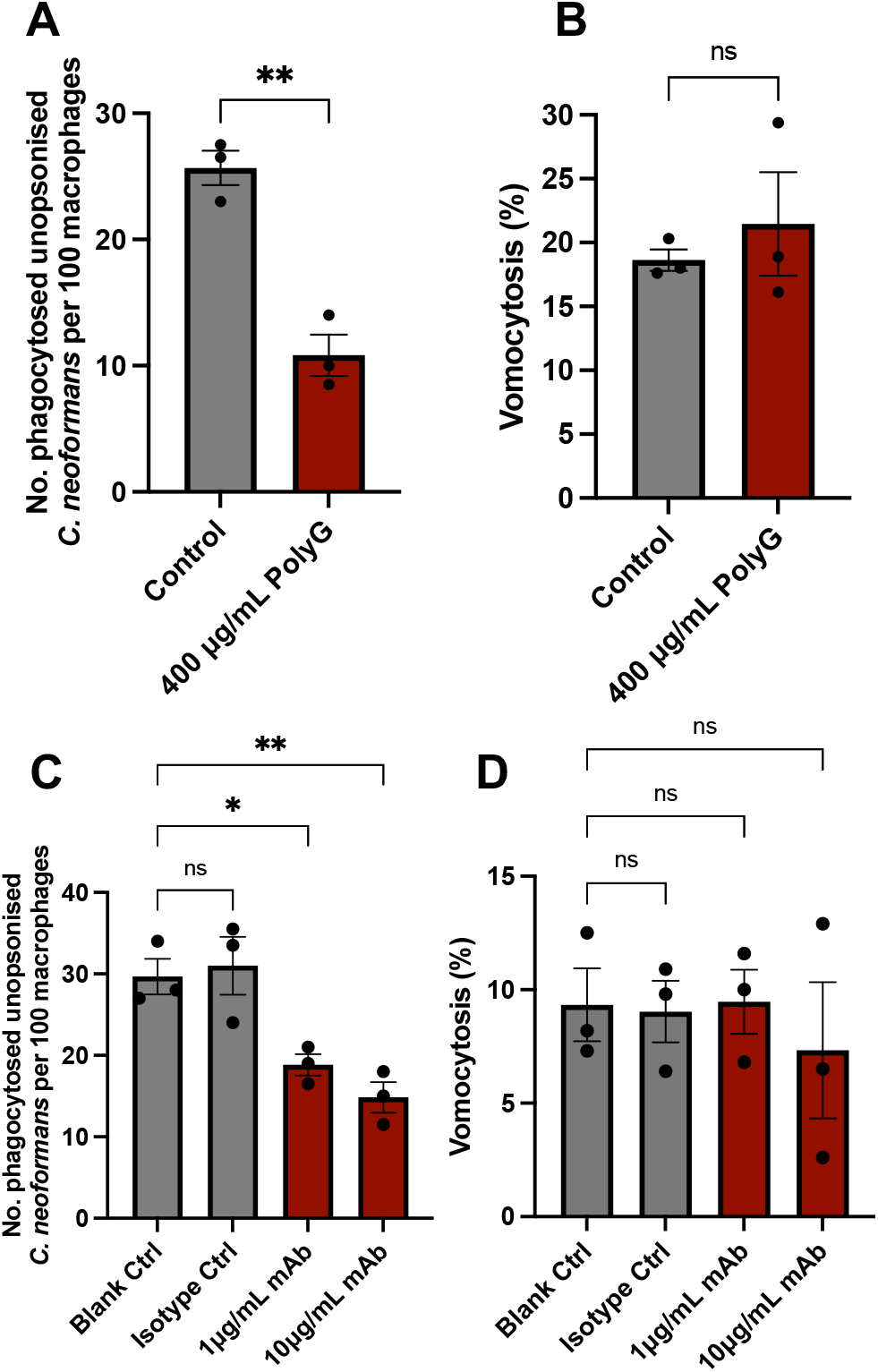
Wildtype macrophages were stimulated overnight with 10 ng/mL LPS. The following day, cells were pre-treated with polyguanylic acid (polyG) **(A, B)** or anti-MARCO ED31 monoclonal antibody (mAb) for 30 mins **(C, D)** then infected with *C. neoformans* still in the presence of polyG or anti-MARCO mAb. Images were acquired every 5 mins for 16 h. **(A, C)** The number of internalised fungi at the beginning of the timelapse video was quantified. **(B, D)** Vomocytosis (%) is the percentage of infected macrophages that experienced at least one vomocytosis events. At least 200 macrophages were quantified per condition. Data is representative of two independent experiments. Data shown as mean ± SEM; ns, not significant; *, p<0.5; **p<0.01 in an unpaired two-sided t-test (**A, B)** and a one-way ANOVA followed by Tukey’s post-hoc test **(C, D)**.

Next, MARCO receptor on wildtype macrophages was blocked using increasing concentrations of an anti-MARCO ED31 antibody. Anti-MARCO antibody reduced MARCO-mediated phagocytosis in a dose-dependent manner (Figure 3C), without impacting the rate of vomocytosis in wildtype macrophages (Figure 3D). According to the manufacturers, the anti-MARCO ED31 antibody recognises the ligand binding domain of MARCO receptors and can therefore compete for receptor binding with *C. neoformans* without impacting intracellular MARCO signalling (24). Taken together, the role of MARCO in vomocytosis is most likely independent of its role in uptake, hence the inability of inhibitors that act on the ligand binding site to phenocopy the genetic knock out of MARCO receptor.

### There is a noticeable difference in actin morphology wildtype and *Marco*^*-/-*^macrophages

Granucci et al. (16) identified a role for MARCO in cytoskeletal remodelling of microglial and dendritic cells. Moreover, repeated actin polymerization and depolymerisation around phagosomes containing cryptococci leading to the formation of transient actin ‘cages’ has been shown to prevent vomocytosis (10). We therefore wondered whether the actin cytoskeleton may be perturbed in MARCO-deficient macrophages.

Rhodamine-conjugated phalloidin staining of uninfected macrophages revealed wildtype macrophages to be more compact than MARCO-deficient macrophages, which were larger and with expansive ruffle-like structures (Figure 4A, white arrows; Supplementary Figure 2). Similarly, in *C. neoformans* infected macrophages, wildtype macrophages appeared more rounded and had well-formed filopodial protrusions (Figure 4B; yellow arrows). Though *Marco*^*-/-*^ cells also had instances of filopodial protrusions from the cell periphery (yellow arrows), these macrophages appeared larger, were less organised and had extensive ruffles (Figure 4B, white arrows). Taken together, there is a clear difference in actin organisation between wildtype and *Marco*^*-/-*^ cells. It is thus possible that MARCO’s role in actin remodelling and organisation is one reason for the elevated vomocytosis seen in *Marco*^*-/-*^ macrophages.

**Figure 4:**
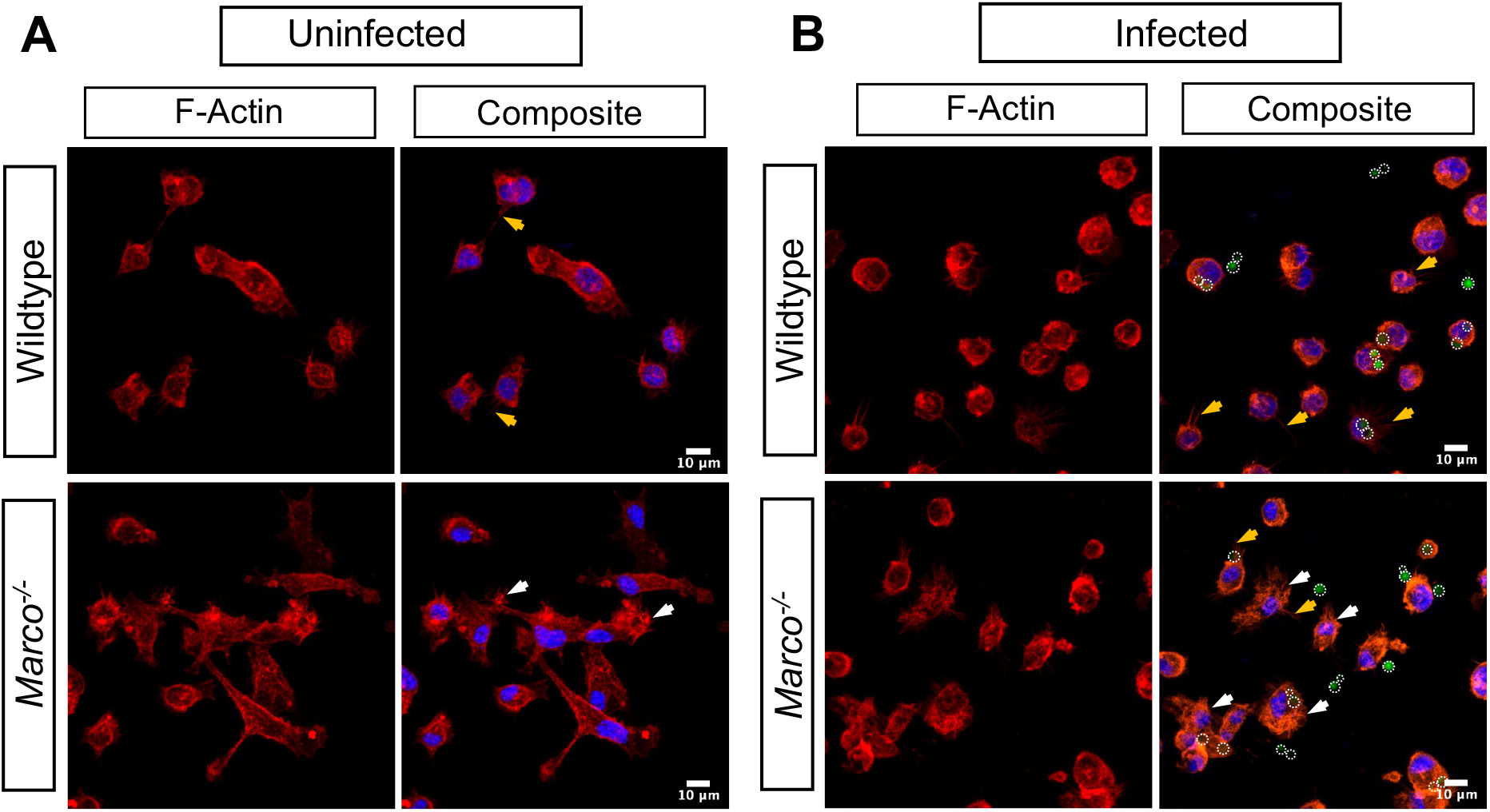
Wildtype and *Marco*^*-/-*^ macrophages were stimulated with 10 ng/mL LPS overnight, left uninfected **(A)** or infected with GFP expressing *C. neoformans* **(B)**. Prior to confocal microscopy imaging, macrophages were fixed, permeabilised and F-actin was stained with rhodamine-conjugated phalloidin. Cells were counter-stained with DAPI to visualize the nucleus, then mounted onto glass slides using Fluoromount mounting medium. Z-stack images were acquired using the Zeiss LSM880 using 63X Oil magnification. Z-stack maximum intensity projection was applied onto the images. Red = F-actin (Phalloidin); Blue = Nucleus; Green with white dashed circle = *C. neoformans*. White arrows show examples of macrophages with ruffle-like structures; Yellow arrows show examples of filopodial protrusions. Scale bar = 10 μm

Our observation raises a number of questions for future investigation. Firstly, given that loss of MARCO leads to elevated vomocytosis, then one role for MARCO may be to sense phagosomal content and prevent premature expulsion, potentially by regulating the formation of actin ‘cages’ the have been show to block phagosome fusion with the plasma membrane (10). It is also possible that MARCO activity is linked to the MAPK ERK5, since ERK5 activity has been implicated in disruptions in actin cytoskeleton during oncogenic transformation (25,26) and is known to modulate vomocytosis (13). Additionally, MARCO may be upstream of Annexin A2, another host signalling molecule found to modulate vomocytosis (14). Annexin A2 plays a significant role in a range of cellular processes including exocytosis and binding to actin to modulate cytoskeleton arrangement (27,28), processes that have been linked to nonlytic expulsion. Finally, we note that this hitherto undocumented impact of MARCO loss on pathogen expulsion will be important for investigators to consider when using *Marco*^*-/-*^ cells or animals for a range of other infection assays.

## Conclusion

Here we present a novel role for MARCO in modulating the vomocytosis of *C. neoformans*. The increase in vomocytosis observed in *Marco*^*-/-*^ macrophages is the most dramatic change in vomocytosis rate observed to date. Increased vomocytosis in *Marco*^*-/-*^ macrophage was also observed when macrophages were infected with a yeast-locked *C. albicans* strain, suggesting that MARCO’s modulation of vomocytosis is a broadly relevant phenomenon. Given that MARCO-deficiency still resulted in elevated vomocytosis of antibody-opsonised *C*. *neoformans* and *C. albicans*, we propose that MARCO’s role in vomocytosis is independent of the mode of uptake, and instead that MARCO may modulate vomocytosis through its role in actin remodelling. We hope this finding will inspire new research aimed at understanding the mechanism and clinical consequence of vomocytosis during host-pathogen interaction.

## Materials & Methods

### Max Plank Institute (MPI) Cell Culture

Max Plank Institute (MPI) cells are a non-transformed, granulocyte-macrophage colony-stimulating factor (GM-CSF)-dependent murine macrophage cell line that is functionally similar to alveolar macrophages (17,29). MPI cell lines isolated from wildtype and MAcrophage Receptor with COllagenous structure knockout (*Marco*^*-/-*^) mice were cultured in Roswell Park Memorial Institute (RPMI) 1640 medium [ThermoFisher] supplemented with 10% heat inactivated FBS [Sigma-Aldrich], 2 mM L-glutamine [Sigma-Aldrich], and 1% Penicillin and Streptomycin solution [Sigma-Aldrich] at 37°C and 5% CO_2_. Each flask was further supplemented with 1% vol/vol GM-CSF conditioned RPMI media prepared using a X-63-GMCSF cell line.

### Phagocytosis Assay

Twenty-four hours before the start of the phagocytosis assay, MPI cells were seeded onto 24-well plates at a density of 2×10^5^ cells/mL in complete culture media supplemented with 1% vol/vol GM-CSF. The cells were then incubated overnight at 37°C and 5% CO_2_. The following day, macrophages were stimulated with 10 ng/mL lipopolysaccharide (LPS) from *Escherichia coli* [Sigma-Aldrich; Cat#: L6529] and 1% vol/vol GM-CSF for 24 h. At the same time, an overnight culture of *Cryptococcus neoformans* var. *grubii* KN99α strain, that had previously been biolistically transformed to express green fluorescent protein (GFP)(30), was set up by picking a fungal colony from YPD agar plates (50 g/L YPD broth powder [Sigma-Aldrich], 2% Agar [MP Biomedical]) and resuspending in 3 mL liquid YPD broth (50 g/L YPD broth powder [Sigma-Aldrich]). The culture was then incubated at 25°C overnight under constant rotation (20rpm).

After overnight LPS stimulation, macrophages were infected with non-opsonised *C. neoformans*. To prepare *C. neoformans* for infection, an overnight *C. neoformans* culture was washed two times in 1X PBS, counted using a haemocytometer, and fungi incubated with macrophages at a multiplicity of infection (MOI) of 10:1. The infection was allowed to proceed for 2 h at 37°C and 5% CO_2_. Where applicable, macrophages were pre-treated with 400 ug/mL polyguanylic acid potassium salt (polyG) [Sigma-Aldrich; Cat#: P4404], rat anti-mouse MARCO ED31 clone monoclonal antibody [BioRad; Cat#: MCA1849], or anti-rat IgG1 isotype control [Invitrogen; Ca#: 14430182] for 30 mins at 37°C prior to infected with non-opsonised *C. neoformans*. In these cases, infection was carried out still in the presence of polyG ligand or antibodies.

For infection with antibody-opsonised *C. neoformans*, 1×10^6^ yeast cells in 100 μL PBS were opsonized for 1 h using 10 μg/mL anti-capsular 18B7 antibody (a kind gift from Arturo Casadevall, Albert Einstein College of Medicine, New York, NY, USA). For infection with heat-killed *C. neoformans*, fungi were killed by heating at 56°C for 30 mins. After 2 h infection at 37°C, macrophages were washed 4 times with PBS to remove as much extracellular *C. neoformans* as possible.

To explore vomocytosis of *Candida albicans*, a yeast-locked *C. albicans* strain was used (a kind gift from Hung-Ji Tsai, University of Birmingham, Birmingham, United Kingdom). The yeast-locked TetOn-NRG1 *C. albicans* strain constitutively express the Nrg1 transcription factor, thereby preventing yeast to hypha transition (18). A colony was suspended in 10 mL YPD broth and incubated overnight at 30°C and 180 rpm. Prior to their use in infection, a *C. albicans* overnight culture was diluted 1:100 in fresh YPD broth, then incubated at 30°C and 180 rpm for 3 h till cells were in exponential phase. Cells in exponential phase were washed with PBS, counted and macrophages were infected at MOI 2:1.

### Time-lapse Imaging

After infection, extracellular fungi were removed and fresh media containing 1% vol/vol GM-CSF (with or without relevant inhibitor) was added back into the wells. For infection with *C. albicans*, imaging began immediately following infection. Live-cell imaging was performed using a Zeiss Axio Observer [Zeiss Microscopy] or Nikon Eclipse Ti [Nikon] at 20X magnification. Images were acquired every 5 mins for 16 hours at 37°C and 5% CO_2_. The resulting videos were analysed using Fiji [ImageJ], at least 200 macrophages were observed, and vomocytosis was scored according to the following guideline:

1. One vomocytosis event is the expulsion of internalized cryptococci from an infected macrophage, regardless of the number of cryptococci expelled if they do so simultaneously.
2. Vomocytosis events are scored as independent phenomena if they occur in different frames or from different macrophages.
3. Vomocytosis events are discounted if the host macrophage subsequently undergoes lysis or apoptosis within 30 min.

### F-Actin Staining and Confocal Microscopy

Macrophages were seeded on 13 mm cover slips placed into 24-well plates. Staining was performed on macrophages fixed with 4% paraformaldehyde for 10 mins at room temperature and permeabilised with 0.1% Triton X-100 diluted in PBS for 10 mins at room temperature. F-actin filaments were stained using 2 units of rhodamine-conjugated phalloidin stain [Invitrogen; Cat#: R415] diluted in 400 μl 1% BSA in 1X PBS and incubated for 20 mins at room temperature. Cells were washed with PBS, then counter stained with 0.5 μg/mL DAPI for 5 mins at room temperature to visualize the nucleus. After PBS washes, glass slides were mounted using Fluoromount mounting medium [Sigma; Cat#: F4680]. Z-stack images were acquired using the Zeiss LSM880 Confocal with Airyscan2, laser lines 405, 488, 561 and 640 nm, and at 63X oil magnification. Image acquisition was performed using the ZEN Black software [Zeiss Microscopy] and the resulting images were analysed using the Fiji image processing software [ImageJ].

### Statistics

GraphPad Prism Version 9 for Mac (GraphPad Software, San Diego, CA) was used to generate graphical representations of experimental data. Violin plots were generated using R programming. Inferential statistical tests were performed using Prism. The data sets were assumed to be normally distributed based on results of Shapiro-Wilk test for normality. Consequently, to compare the means between treatments, the following parametric tests were performed: unpaired two sided t-test, one-way ANOVA followed by Tukey’s post-hoc test. When data failed the normality test, Mann-Whitney U nonparametric test was used. Variation between treatments was considered statistically significant if p-value < 0.05.

## Acknowledgments

We thank Hung-Ji Tsai for the provision of yeast-locked *C. albicans* strain. C.U.O is supported by a PhD studentship from the Darwin Trust of Edinburgh. R.C.M. gratefully acknowledges support from the BBSRC and European Research Council Consolidator Award.

---

**Supplementary Figure 1:**
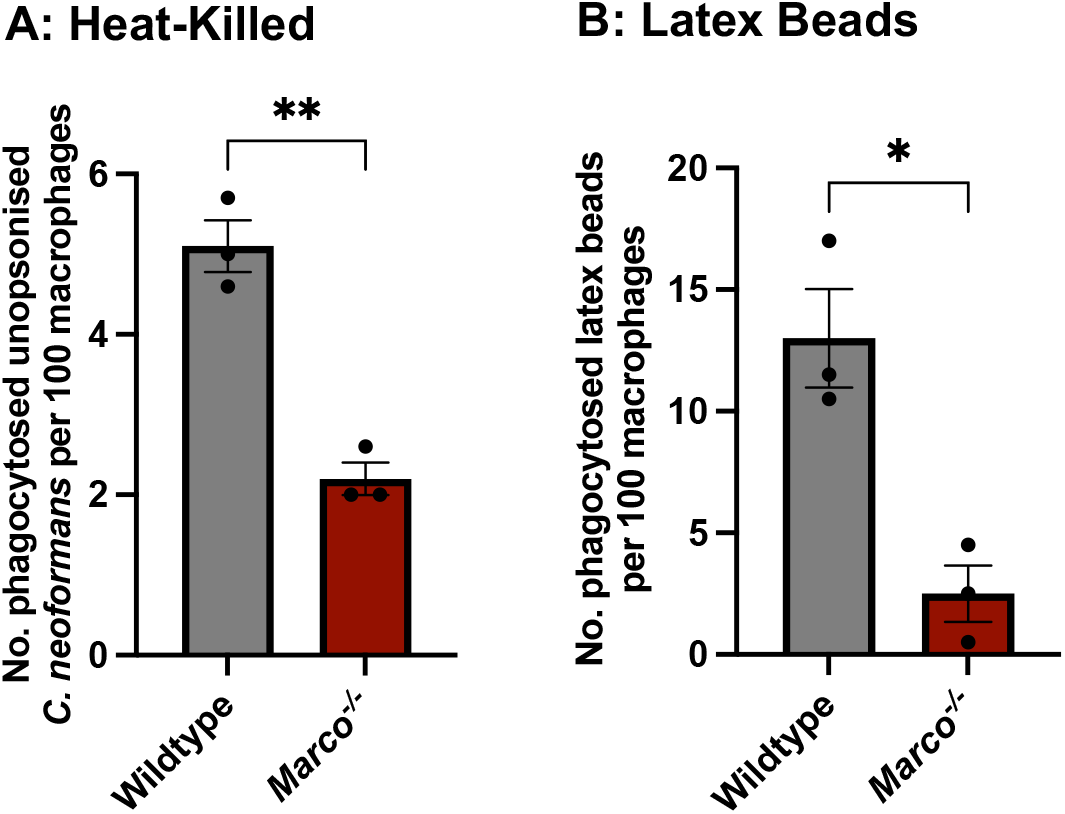
Wildtype and *Marco*^*-/-*^ macrophages were infected with **(A)** *C. neoformans* killed by heating at 56°C for 30 mins or **(B)** 7 μm latex beads. After 2 h infection, images were acquired every 5 mins for 16 h. The number of internalised cryptococci or latex bead per 100 macrophages was quantified. At least 200 macrophages were observed per condition. Data shown as mean ± SEM; *p<0.05; **p<0.01 in a t-test. Data is representative of two independent experiments.

**Supplementary Figure 2:**
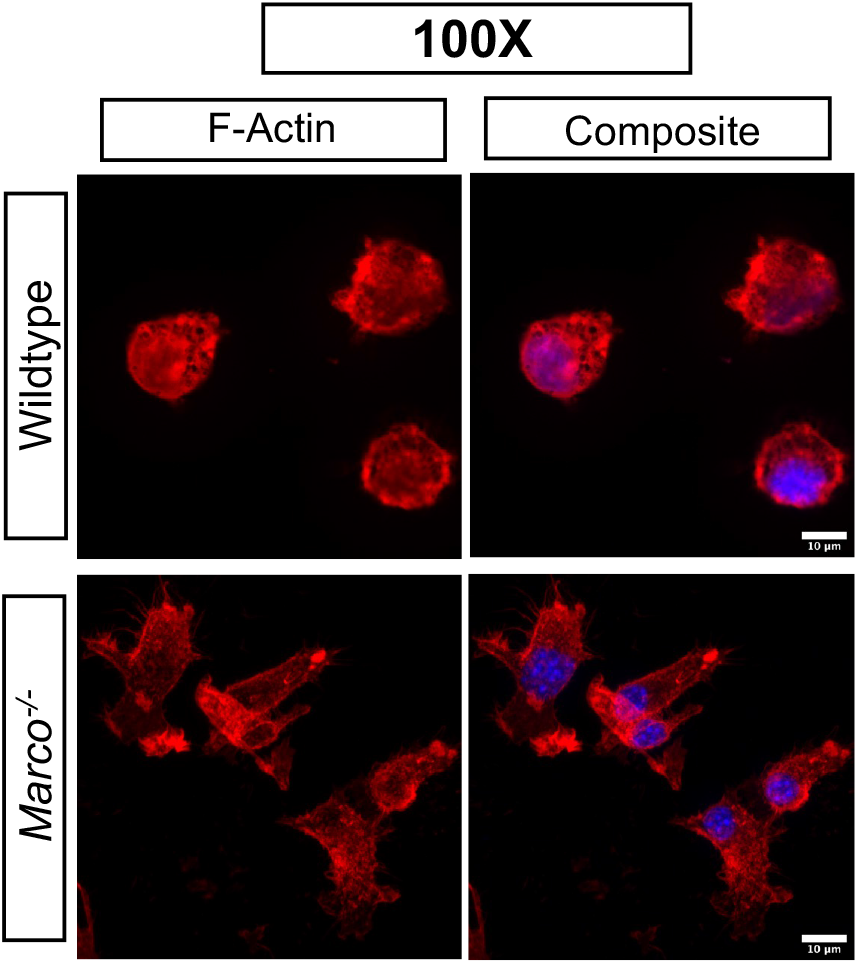
Wildtype and *Marco*^*-/-*^ macrophages were stimulated with 10 ng/mL LPS overnight, then stained with rhodamine-conjugated phalloidin to visualize F-actin distribution. Z-stack images were acquired on the Zeiss LSM880 using 100X Oil magnification. Z-stack maximum intensity projection was applied onto the images. Red = F-actin (Phalloidin); Blue = Nucleus. Scale bar = 10 μm

**Supplementary Video 1:** Representative video showing vomocytosis of *C. neoformans* from *Marco*^*-/-*^macrophages. Video corresponds to Figure 1D. Time shown as: hh:mm:ss

**Supplementary Video 2:** Example video showing rapid time-to-vomocytosis in *Marco*^*-/-*^macrophages. Time shown as: hh:mm:ss

**Supplementary Video 3:** Representative video showing vomocytosis of *C. albicans* from *Marco*^*-/-*^macrophages. Video corresponds to Figure 2E. Time shown as: hh:mm:ss

## Notes

### Competing Interest Statement

The authors have declared no competing interest.

